# Genome-scale transcriptome augmentation during *Arabidopsis thaliana* photomorphogenesis

**DOI:** 10.1101/2025.01.30.635720

**Authors:** Geoffrey Schivre, Léa Wolff, Filippo Maria Mirasole, Elodie Armanet, Mhairi L. H. Davidson, Adrien Vidal, Delphine Cuménal, Marie Dumont, Mickael Bourge, Célia Baroux, Clara Bourbousse, Fredy Barneche

**Author notes:** & (Clara Bourbousse, Fredy Barneche).

## Abstract

Plant photomorphogenesis is a light-induced developmental switch that combines massive reprogramming of gene expression and a general enhancement in RNA Polymerase II activity. Yet, transcriptome analyses have failed to demonstrate a clear tendency toward gene upregulation. To solve this conundrum, we optimized a spike-in RNA-seq experimental and analysis pipeline and reconciled transcriptome dynamics with epigenomic and cytogenetic observations. We found that light induces a two-fold expansion of the transcriptome during *Arabidopsis thaliana* cotyledon photomorphogenesis, with 94% of the 9,128 differentially expressed genes being upregulated within the first six hours. Transcriptome augmentation was further detected at a similar strength in spike-free RNA-seq datasets from independent laboratories renormalized using stable genes that mimic the spike-in information. On this new basis, reinterpretation of light-mediated gene regulatory pathways revealed a quasi-exclusive positive effect of HY5 and other key light-induced transcription factors at target genes. This new standpoint unveiled the unilateral impact of light on the transcriptome of Arabidopsis cotyledons at the genome scale and opens new avenues for investigating global genome regulation during plant developmental and environmental responses.

---

Plants display extraordinary developmental plasticity to external cues, especially when it comes to light. In addition to fueling photosynthesis, light is sensed and signaled to continuously modulate plant physiology, development, and cellular status. These dynamic adjustments to local environmental conditions are achieved largely through the regulation of gene expression patterns^1–3^. The spectacular effect of light on plant gene expression has been mostly studied when a germinating plant is exposed to light for the first time, thereby triggering de-etiolation, the transition from skoto-to photomorphogenesis. In complete darkness, DEETIOLATED-1 (DET1), CONSTITUTIVE-PHOTOMORPHOGENIC-1 (COP1), and several other repressors of photomorphogenesis induce seedling etiolation by promoting hypocotyl (stem) elongation and inhibiting the development of embryonic leaves (cotyledons)^4–6^. Once red, blue, or UV-B light is perceived by photoreceptors, complex signaling transduction events induce photomorphogenesis with the onset of chloroplast biogenesis and photosynthesis^3,7^. Former studies indicated that as much as 40% of the nuclear genes are differentially expressed during the transition^8–12^ in an organ- and cell-specific manner through the action of multiple light-regulated transcription factors (TF) such as PHYTOCHROME INTERACTING FACTORS (PIFs), HYPOCOTYL ELONGATION-5 (HY5) and TEOSINTE BRANCHED1/CYCLOIDEA/PCF (TCPs)^8–11,13^

In cotyledons, light-mediated transcriptome changes involve gene-specific and large-scale variations in chromatin composition and 3D organization^14,15^ together with nucleus expansion and endoreduplication, a mechanism by which the nuclear DNA content increases^11,16^. This general adaptation of the nucleus and epigenome may sustain a global increase in transcription, evidenced by a higher proportion of RNA Polymerase II (RNA Pol II) engaged in transcript production^17^ and an enrichment at most genes of monoubiquitinated histone H2B (H2Bub)^18,19^, a histone modification functionally associated with RNA Pol II elongation. Induction and maintenance of a hypotranscriptional status in darkness may result from a genome-wide attenuation of transcription, which would be released upon light perception through a massive wave of gene induction. This initial evidence points to an augmentation of global cellular mRNA levels in response to light. However, transcriptomic investigations have consistently identified approximately the same number of genes being up- or down-regulated during seedling or cotyledon deetiolation, and none has demonstrated a clear trend in gene upregulation^8–11^. This conundrum most plausibly results from the capacity of former transcriptomic analyses to report relative changes in transcript abundance while ignoring the fundamental importance of transcript absolute quantification. Multiple conceptual and experimental studies have identified the technical difficulties in quantifying absolute variations in mRNA steady-state levels using RNA-seq, especially when cellular transcript abundance is different between samples^20–24^. Using standard bulk or single-cell RNA-seq methodologies, the information needed to estimate absolute transcript abundance is usually lost during sample collection or RNA extraction and further vanishes upon data normalization, count data generated by sequencing technologies being compositional^25^. Hence, while former studies gained insightful information on light-mediated gene expression changes during photomorphogenesis and other developmental transitions, we ignore how transcription intensification impacts transcriptome dynamics in plant cells. In addition, we also lack estimates to properly integrate transcriptome data with genomic profiling of RNA Pol II, nascent transcripts, transcription-associated factors, or epigenome changes.

In this study, combining epigenomic profiling and live imaging, we first report that light sensing triggers a genome-scale augmentation of the transcriptional regime, marked by a doubling of RNA Pol II activity in individual nuclei. We further reveal how transcription intensification functionally impacts transcriptomic patterns during *A. thaliana* cotyledon photomorphogenesis. To overcome the technical bottlenecks of quantitative RNA-seq approaches, we set up a dedicated spike-in RNA-seq experimental and analysis pipeline. This showed that cotyledon photomorphogenesis leads to 94% upregulation *vs* 6% downregulation resulting in a 2-fold increase in transcriptome size. This approach further enabled the identification of stably expressed genes during cotyledon de-etiolation that we used to renormalize publicly available transcriptome datasets, thereby demonstrating the possibility of recovering accurate information on light-mediated transcriptome size augmentation in independent studies lacking a spike-in approach. These results reconcile insights into global changes in steady-state RNA levels with epigenome dynamics and RNA Pol II activity but also provide new information on gene expression regulation by light.

## Results

### Cotyledon photomorphogenesis involves a general increase in RNA Pol II amount and activity

To estimate the levels of RNA Pol II engaged in transcript production in single nuclei from skotomorphogenic versus photomorphogenic cotyledons (**Fig. 1A**), we used an immunocytometry approach employing two antibodies recognizing the elongating Serine 2 phosphorylated (S2P) and the inactive unphosphorylated (NP) RNA Pol II forms. This first showed that both S2P and NP RNA Pol II levels per nucleus increased proportionally to ploidy (**Fig. 1B and Figure S1A**), in agreement with the proposed role of endoreduplication in sustaining high messenger RNA (mRNA) production rates^26–30^. Corroborating previous analyses^17^, we also detected a 1.6-to 2-fold higher proportion of elongating RNA Pol II in cotyledon nuclei of light-grown compared to dark-grown seedlings (**Fig. 1B)**. The effect of light was significant for each ploidy class and therefore independent of the amount of DNA serving as a template for transcription. Last, analyzing each antibody signal separately showed that the S2P form is more abundant in the light than in the dark, while the NP form is less so (**Fig. 1B and Figure S1A)**. This contrast increases with ploidy, suggesting that RNA Pol II is differentially impacted depending on the number of genome copies in the nucleus, the cell type, or the differentiation status.

**Figure 1.**
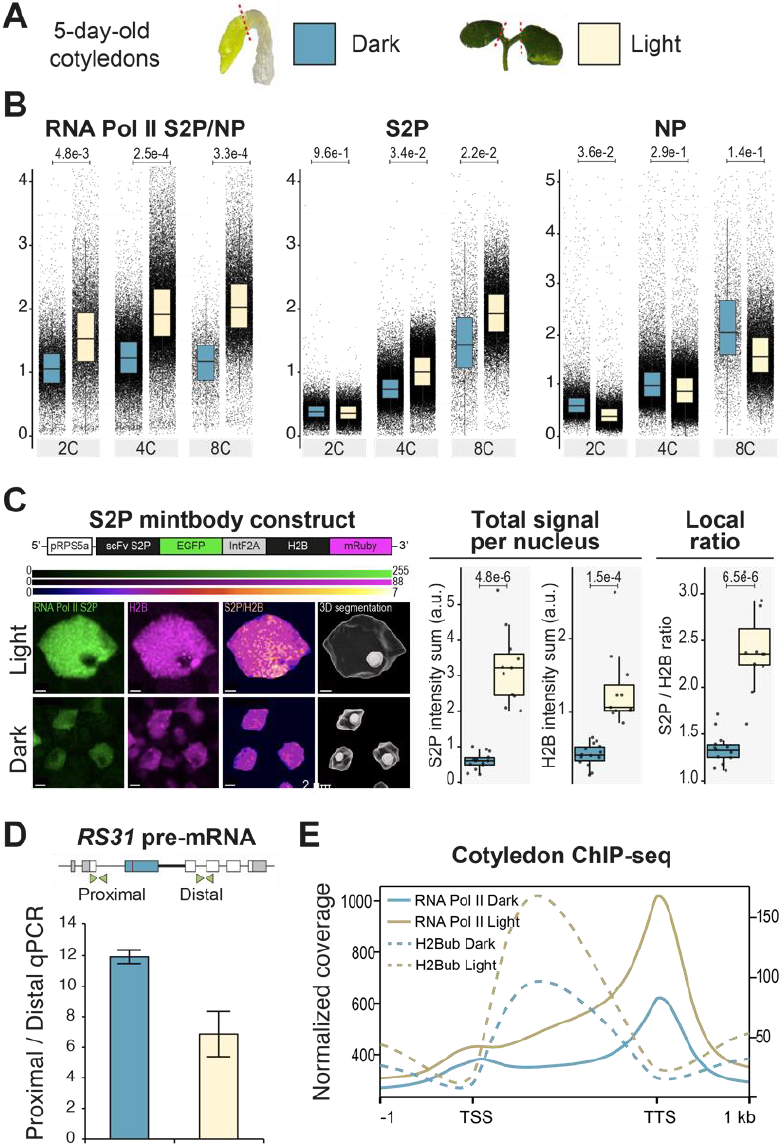
Light impacts the transcriptional status of cotyledon nuclei. **A.** Analyses in this panel employed cotyledons dissected from dark- or light-grown seedlings. **B.** RNA Pol II activity in individual cotyledon nuclei was determined by flow cytometry as the ratio of fluorescent signals after immunolabeling of its Serine 2 phosphorylated (S2P) and non-phosphorylated (NP) forms for each ploidy class. For the ratio, the median of the 2C Dark sample was arbitrarily set to 1. Each dot represents the measure in a single nucleus. Three biological replicates were used. The p-value from a t-test is given between dark and light for each ploidy class. (See Source data 1) **C.** Live imaging of cotyledon mesophyll nuclei expressing RNA Pol II S2P mintbody-GFP and H2B-mRuby reporting local signals of the elongating RNA Pol ll and chromatin. Left panel: 3D projections of deconvolved images showing the levels of RNA Pol II S2P (green), H2B-mRuby (magenta), and their ratio (“Fire”)^31^. See **Figure S1B** and Source data 1 for further details. Right panel: quantification of S2P and H2B total nuclear signals and RNA Pol II-S2P:H2B signal ratio (n=14 and n=11 for dark- and light-grown seedlings, respectively). **D.** RT-qPCR analysis of *AtRS31* pre-mRNA accumulation. The bars indicate the ratio of the signal obtained with primer pairs recognizing a proximal and a distal position on the *AtRS31* pre-mRNA. **E.** ChIP-seq metaprofiles of RNA Pol II S2P and H2Bub over the ensemble of genes marked in Dark or Light samples.

To track RNA Pol II activity in live cotyledons, we imaged a ratiometric Modification-specific INTracellular antiBODY (mintbody) S2P RNA Pol II reporter line^31^ (**Fig. 1C**). Following two-photon 3D microscopy, we quantified S2P mintbody-GFP vs H2B-mRuby live fluorescence levels per pixel in cotyledon nuclei of dark and light-grown seedlings (**Figure S1B**). On average, local S2P signals normalized to chromatin and total S2P RNA Pol II signal in whole nuclei were higher in the light (**Fig. 1C**), showing, *in vivo*, that RNA Pol II activity is induced ∼2-fold during photomorphogenesis independently of DNA content.

We then asked whether light-triggered enrichment in active RNA Pol II associates with differences in RNA Pol II kinetics. We focused on *At-RS31* nascent RNA accumulation at proximal (P) vs distal (D) regions, which can be used as a proxy of low RNA Pol II processivity^32,33^. RT-qPCR analysis of *At-RS31* pre-mRNA detected a higher P/D ratio in dark-grown than in light-grown cotyledons (**Fig. 1D**), indicating a lower progression of RNA Pol II in the dark at this locus.

Finally, to gain a genome-wide view of RNA Pol II activity during cotyledon photomorphogenesis, we profiled H2Bub, a co-transcriptional histone modification, and S2P RNA Pol II in dissected cotyledons by ChIP-seq. In line with a general hypotranscriptional status of skotomorphogenic cotyledon nuclei, both S2P RNA Pol II and H2Bub chromatin levels at gene bodies were lower in dark than in light-grown cotyledons (**Fig. 1E and Figure S2A)**. Collectively, our cytogenetic and epigenomic analyses showed that cotyledon photomorphogenesis involves a general increase of RNA Pol II content and activity over the genome.

### Light-mediated augmentation of the transcriptional regime is not detected using standard RNA-seq

Having established that RNA Pol II activity and transcription increase during cotyledon photomorphogenesis, we questioned to what extent such a genome-scale control would impact the transcriptome patterns. RNA-seq analysis of 200 cotyledons dissected from dark- or light-grown seedlings showed no tendency for a global increase in mean transcript levels or gene expression fold changes (Log_2_ FC distributed around 0, *i.e*., no difference), and only a minor bias in the number of down- or up-regulated genes (43 vs 57%, respectively, with an FDR < 0.01 and a |Log_2_ FC| > 0.5) (**Fig. 2A**). Hence, as in previous studies^10,11^, a standard RNA-seq methodology detected no global difference that would reflect a 2-fold intensification of transcription.

**Figure 2.**
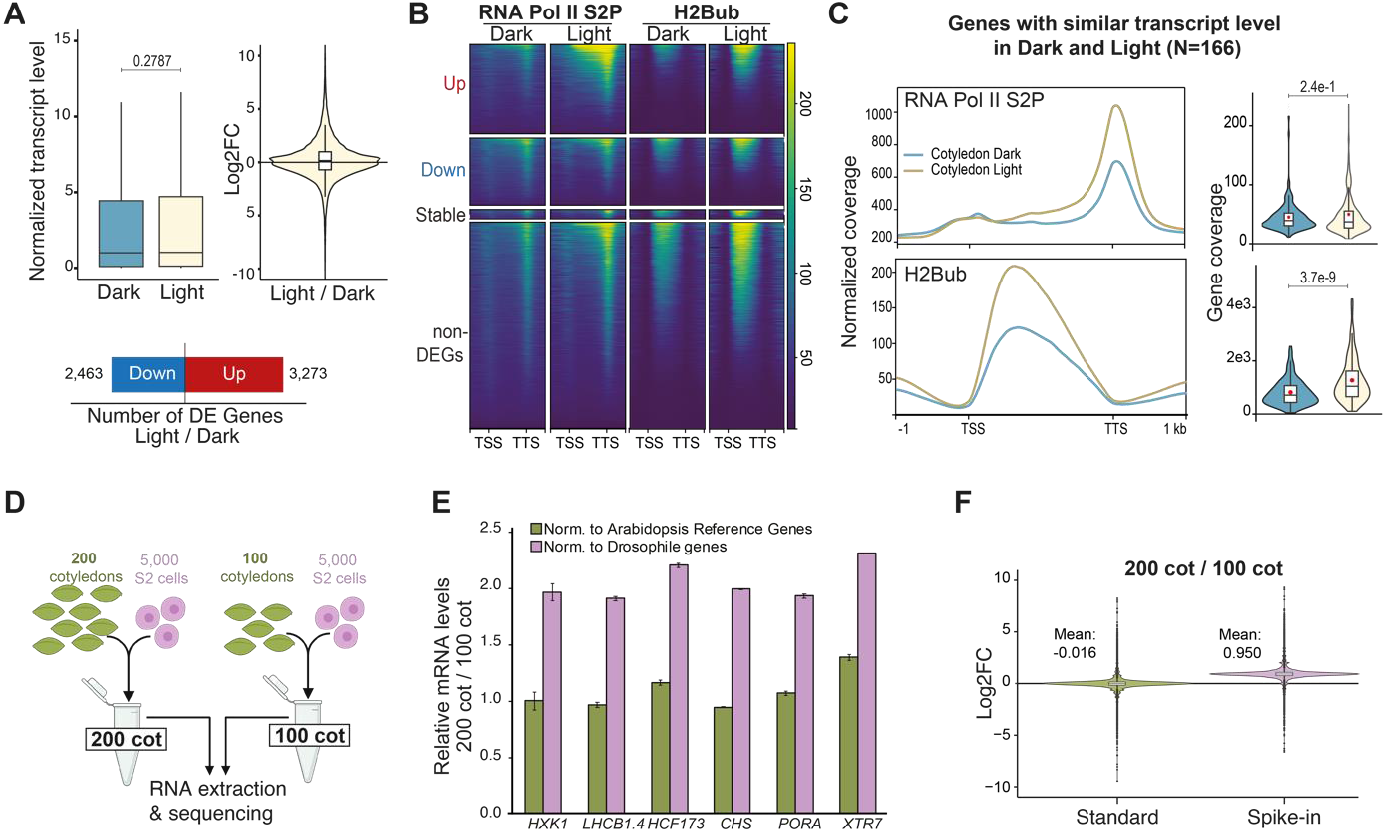
Transcription intensification is not detected using standard RNA-seq. **A.** Distribution of transcript levels, expression fold changes (Log_2_), and numbers of differentially expressed genes (q-value < 0.01 for a |Log2 FC| > 0.5) using a standard RNA-seq analysis of 5 biological replicates comparing cotyledons dissected from dark- or light-grown seedlings. **B.** RNA Pol II S2P and H2Bub chromatin enrichment at the genes analyzed in A. **C.** RNA Pol II S2P and H2Bub meta-profiles over all genes displaying stable expression in standard RNA-seq analysis of dissected cotyledons (q-value < 0.01 for a |Log_2_ FC| < 0.5). **D.** Experimental setup for introducing Drosophila S2 cells as an exogenous spike-in reference (RNA-Rx). As a proof-of-concept experiment, 5,000 S2 cells were mixed with 100 or 200 dissected cotyledons before RNA extraction. **E.** RT-qPCR analysis of mRNAs extracted from 100 or 200 dissected cotyledons, both including the same number of Drosophila cells as a spike-in. The ratio of RNA levels in the two sample types was obtained either by normalizing to Arabidopsis reference genes (*AT3G02065, AT1G13320,* and *AT2G41020*) or Drosophila genes (*RpL21, cg25c, vkg, B4,* and *pum*). **F.** Distribution of expression changes (Log_2_ FC) in four biological replicates. RNA sequencing reads were analyzed by normalizing solely to library size (Standard) or using reads mapping to Drosophila genes (Spike-in normalization).

To assess this discrepancy at the gene level, we compared S2P-RNA Pol II and H2Bub levels at loci encoding differentially or stably expressed genes (FDR < 0.01 and |Log_2_ FC| > 0.5 or |Log_2_ FC| < 0.5, respectively). While light-induced genes displayed higher S2P-RNA Pol II and H2Bub mean enrichment in light than in darkness, down-regulated genes showed very poor or no change of these two transcription hallmarks (**Fig. 2B and Figure S2B**). This suggested that consideration of this gene set as being downregulated may be artifactual. Conversely, the set of stably expressed genes was enriched in S2P-RNA Pol II and H2Bub in the light compared to darkness (**Fig. 2C)**, suggesting that changes in transcript levels were under-estimated. Hence, discrepancies between S2P-RNA Pol II or H2Bub profiles and standard RNA-seq analysis suggested that the effects of transcription intensification were largely missed, possibly underestimating the impact of light on transcript levels during cotyledon photomorphogenesis.

This seeming contradiction most likely resulted from the well-described inability of standard RNA-seq analyses to inform on absolute variations of transcript levels^20,22,23,34^. Indeed, count data generated by sequencing technologies being compositional, a strong limitation of RNA-seq standard normalization procedures is that they estimate changes in transcript levels relative to each other but not absolute changes between samples^25^. To circumvent this issue, we optimized the use of an exogenous reference. Considering that spiking in exogenous RNA before library preparation can improve data normalization but cannot inform on absolute levels in the starting material because of RNA extraction and processing biases (**Figure S3A)**, we spiked a fixed number of *D. melanogaster* S2 cells directly in cotyledon lysates and extracted *A. thaliana* and *D. melanogaster* RNA together. As a proof of concept, we compared transcript levels in pools of 100 and 200 cotyledons dissected from light-grown siblings, meant to display a doubling of all transcript levels when normalizing to the spike-in reference (**Fig. 2D**). While standard RT-qPCR normalization using housekeeping genes expectedly gave similar relative transcript levels in both sample types, using Drosophila transcripts as reference allowed the detection of twice more *A. thaliana* transcripts in the “200 cot” than in the “100 cot” samples for all tested genes (**Fig. 2E**). Extending this principle to RNA-seq expectedly enabled the detection of a two-fold shift in the relative proportion of Arabidopsis vs Drosophila reads in all four biological replicates comparing 100 and 200 cotyledons (**Figure S3B)**. To robustly normalize RNA-seq data to the spike-in, we developed an ad hoc analysis pipeline hereafter called RNA-Rx (RNA-seq with reference exogenous transcripts) that mitigates the technical variability induced by the spike-in (See the methods section and Source Data 2) while reducing data dispersion as much as standard RNA-seq procedures (**Figure S3C**). This confirmed a 2-fold change using the RNA-Rx pipeline while no global change could be detected using a standard approach (**Fig. 2F and Figure S3D**). These observations demonstrate that RNA-Rx enables absolute comparisons of transcript levels between samples.

### Average cell transcriptome size increases during cotyledon photomorphogenesis

A pre-requisite for inter-sample comparisons with RNA-Rx is to use samples with similar cell or nucleus numbers. This is roughly the case of cotyledons from dark- and light-grown seedlings^35^. Yet, in the light, about 20% of the cotyledon cells are subjected to one more endoreduplication cycle producing extra genome copies in the light (average ploidy level of 3.5 *vs* 4.3; **Figure S4B**). Conversely, cotyledons display neither cell division nor ploidy changes during the first 24 hours of de-etiolation and therefore are perfectly suited for an RNA-Rx approach (**Figure S4A-B,** López-Juez et al. 2008^11^). We ensured that RNA Pol II processivity increases after 1, 6, and 24 hours of light exposure by determining *At-RS31* pre-mRNA proximal *vs* distal sites. We indeed observed a fast and strong decrease in the P/D ratio, reaching 2-fold after 1 hour (**Figure S4C**). Consequently, in subsequent analyses, dark-grown seedlings were exposed to continuous light for 1 to 24 hours before cotyledon dissection (**Fig. 3A**).

**Figure 3.**
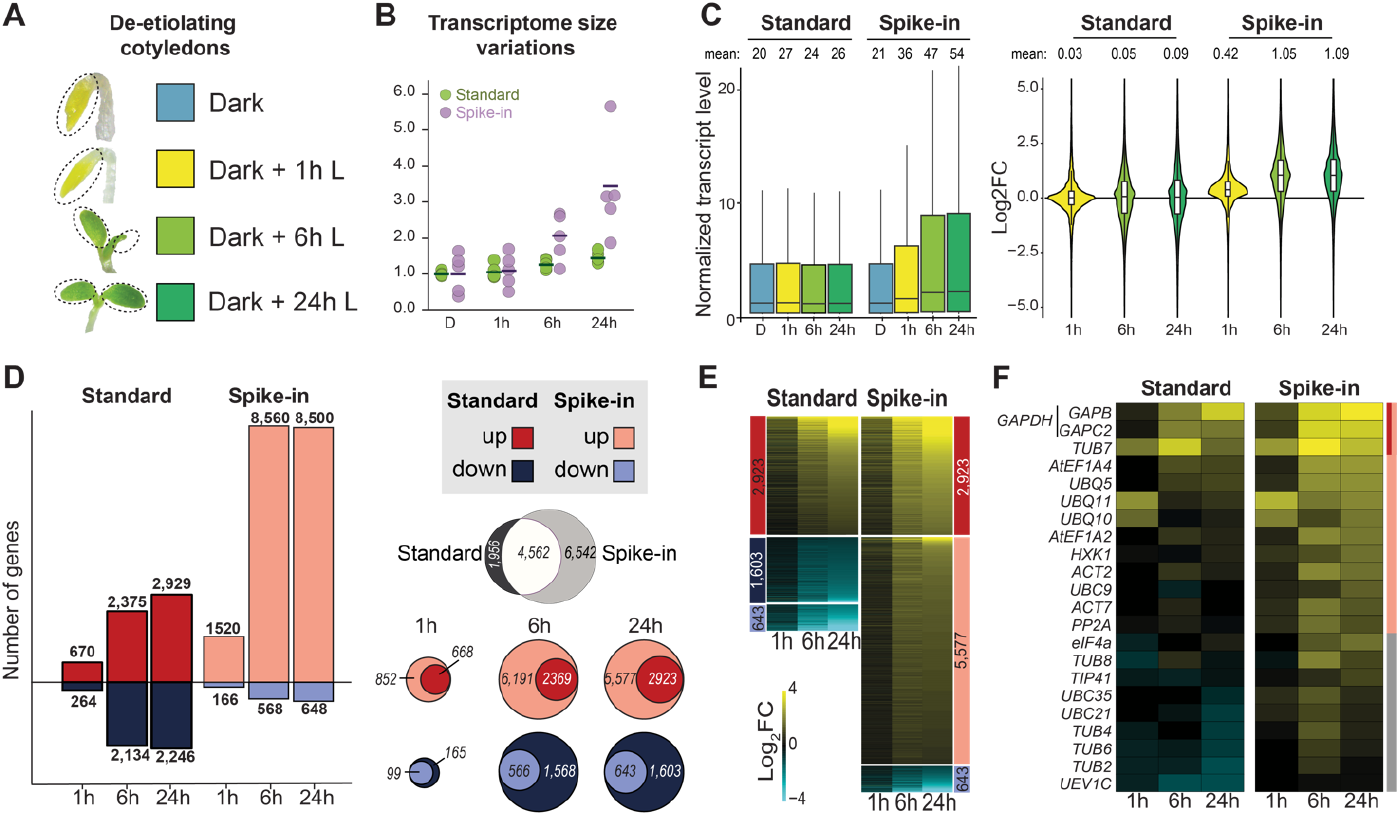
Light induces transcriptome size increase in cotyledon cells during de-etiolation. **A.** The analyses employed cotyledons dissected from 5-day-old seedlings grown in darkness (Dark) or exposed to light for the indicated duration. **B.** Transcriptome size of the five biological replicates was estimated from the size factor of DESeq2 differential analyses of Standard RNA-seq and Spike-in RNA-Rx divided by library size. For comparison, the median values of the Dark samples were arbitrarily set to 1. **C.** Distribution of mean transcript levels (left panel) and Log_2_ FC of expression (right panel). The Dark time point was used as a reference for each Standard RNA-seq and Spike-in RNA-Rx analysis. **D.** Number of differentially expressed genes (DEGs, q-value < 0.01 for a |Log2 FC| > 0.5) in standard RNA-seq and Spike-in RNA-Rx analyses. The Venn diagrams show the number of DEGs commonly called in the two methods. **E.** Log_2_ FC of expression at three time points of light exposure compared to the Dark time point for all genes analyzed in D. Genes are ranked according to their Log_2_ FC at the 24 h time point. **F.** Heatmap representing the Log_2_ FC of expression at the indicated time points compared to the Dark time point for genes ordinarily used as RT-qPCR references or considered as housekeeping. Genes are ranked according to the expression Log_2_ FC at the 24-hour time point. The sidebar indicates when gene sets were called DEGs in Standard RNA-seq or Spike-in RNA-Rx analyses following the color code displayed in D.

We used this setup for an RNA-Rx time series analysis of cotyledon photomorphogenesis. An Arabidopsis cotyledon being made of ca. 4,000 cells^36^, the number of cells in each sample was estimated at ∼800,000 irrespective of the light condition, to which we added 5,000 Drosophila S2 cells (0.625%) per sample in five independent biological replicates. Dual mapping of the sequencing reads to the Drosophila and Arabidopsis reference genomes first showed that the percentage of Drosophila *vs* Arabidopsis reads decreased with light exposure from an average of 2.26% at time 0 (T0 Dark) to 0.81% after 24 hours (**Figure S5A**), indicating a progressive augmentation of cells’ transcriptome sizes during de-etiolation (**Fig. 3B**). The distribution of transcript levels and gene expression fold changes were expectedly stable in response to light using standard normalization, but increased over time with spike-in normalization, reaching a 2-fold rise after 6 hours (**Fig. 3C, Figure S5B and E**). Both observations support an average doubling of transcriptome size in cotyledon cells during photomorphogenesis.

As anticipated, transcript level variations were more dispersed for cotyledon photomorphogenesis than when comparing 100 *vs* 200 cotyledons, reflecting the extensive gene reprogramming events in addition to transcriptome size doubling (**Fig. 3C and Figure S5B**). Compared to standard RNA-seq, RNA-Rx identified almost twice as many differentially expressed genes at one or more time points, affecting 46% of the total number of Arabidopsis genes (11,104 among the 23,898 genes analyzed) (**Fig. 3D**). Last, while standard RNA-seq detected comparable numbers of down- and upregulated genes as in former studies^9–11,13^, RNA-Rx identified a considerable bias toward gene upregulation with 8,500 genes upregulated but only 648 genes downregulated after 24 hours (**Fig. 3D, Figure S5B**). Accordingly, about 6,000 genes were found upregulated only when using the spike-in normalization. Conversely, a large majority of the genes considered downregulated with standard RNA-seq were not differentially expressed according to the RNA-Rx analysis (**Fig. 3D-E, Figure S5D**). In agreement with our observation of greater amounts of RNA Pol II in individual nuclei (**Fig. 1**), RNA-Rx identified genes encoding RNA Pol II subunits as being upregulated (**Figure S5C**). RNA-Rx also identified many genes commonly used as references in expression studies, such as *ACT2, UBQ10, PP2A,* and *GAPDH*, as upregulated (**Fig. 3F**), indicating that housekeeping genes are part of the transcriptome augmentation process. Overall, RNA-seq with spike-in followed by *ad hoc* normalization revealed that cotyledon de-etiolation involves a wide increase in transcript levels formerly masked when using standard methodologies, while hundreds of genes formerly seen as downregulated are actually stable.

### Reinterpreting the mechanistic and functional gene regulatory paths of cotyledon photomorphogenesis

Considering that switching from standard to spike-in normalization may impact current views on light-driven gene regulation, we examined the expression of genes targeted by the major transcription factors mediating light signaling. Compared to standard RNA-seq, a much higher fraction of the genes specifically bound in the light by HY5, GLK1/2, FHY1, BZR1, TCP4, or GI were induced during cotyledon de-etiolation. For example, RNA-Rx identified that, among the HY5 target genes differentially expressed during the transition, 93% (185/198) but not 66% (90/135) are upregulated (**Fig. 4A-B)**. RNA-Rx identified fewer downregulated genes in response to light than standard RNA-seq, the most striking example being GLK1/2 target genes displaying 1.6% instead of 7% downregulated target genes after 24 hours. Hence, RNA-Rx almost exclusively identified a GLK1/2 and HY5 propensity for gene activation, better supporting functional analyses^13,37,38^. For dark-activated TFs such as PIF3, FHY3, ARF6, and BZR1, RNA-Rx also detected more light-induced genes and less light-repressed genes than standard RNA-seq. In the case of PIF3 target genes, RNA-Rx detected 240 instead of 83 upregulated genes and 145 instead of 265 downregulated genes after 24 hours of light (**Fig. 4A-B**), in agreement with PIF3 acting both as a transcriptional activator and repressor in etiolated cotyledons^39–41^.

**Figure 4.**
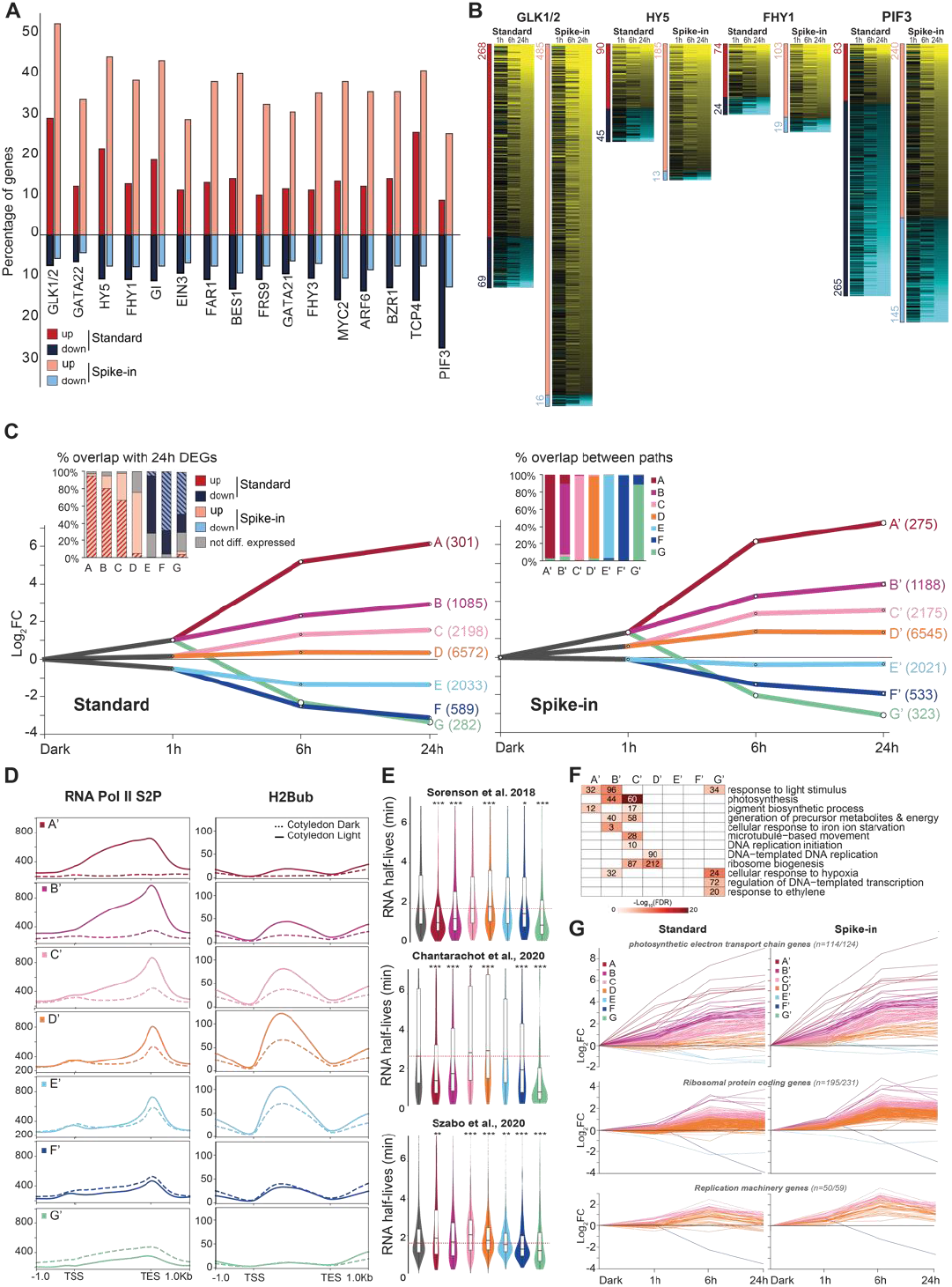
RNA-Rx enables revisiting cotyledon de-etiolation gene regulatory paths. **A.** Percentage of genes up- or downregulated after 24 h of light (q-value < 0.01 for a |Log2 FC| > 0.5) amongst transcription factor target genes. **B.** Expression Log2 FC of genes bound by GLK1/2, HY5, FHY1, and PIF3 transcription factors and differentially expressed upon 24 h of light exposure using Standard or Spike-in RNA-Rx analyses, respectively. Genes are ranked according to the Log2 FC at 24 h. **C.** DREM analysis of coregulated gene paths using expression Log_2_ FC from Standard RNA-seq and Spike-in RNA-Rx. Seven paths were found in both cases, called A to G (Standard) or A’ to G’ (Spike-in) that largely overlap. The left inlay shows the percentage of DEGs defined using Standard RNA-seq or Spike-in RNA-Rx in the different paths at the 24 h time point. The right inlay shows the percentage of genes found commonly in each path after Standard RNA-seq and Spike-in RNA-Rx. **D.** Metaplot profiles of RNA Pol II S2P and H2Bub enrichment in dark and light-grown cotyledons at the genes corresponding to the DREM paths A’ to G’. **E.** Distribution of mRNA half-lives in the DREM paths A’ to G’ according to Sorenson et al. (2018), Chantarachot et al. (2020), and Szabo et al. (2020)^43–45^. The distributions over all TAIR10 genes are shown as reference in gray and compared to each DREM path with a student t-test (* < 0.1, ** < 0.01 and ***< 0.001). **F.** Over-represented gene ontology (GO) categories within the DREM path A’ to G’. Numbers correspond to the genes falling in each GO category. **G.** Variations of expression Log_2_ FC upon Standard RNA-seq or Spike-in RNA-Rx analysis of genes encoding proteins of the photosynthetic chain, ribosomes, and replication machinery.

We then compared the regulatory networks inferred from RNA-Rx *vs* standard RNA-seq using Dynamic Regulatory Events Miner (DREM)^42^. Employing identical gene sets and parameters, the two analyses identified seven modules of expression with similar gene sets but different patterns (**Fig. 4C, Figure S6A**). Genes that were highly induced in response to light displayed an upregulation twice as strong using RNA-Rx (paths A’-C’) than standard RNA-seq (paths A-C) data (**Figure S6A**). Path D and D’ contain most of the genes upregulated in RNA-Rx but not in standard RNA-seq data (75%). Path E and E’ are mainly composed of genes downregulated in standard RNA-seq (68%) but not in RNA-Rx. Finally, paths F-G and F’-G’ correspond to genes downregulated in both analyses but with a milder intensity in RNA-Rx datasets.

To test the validity of the gene expression paths defined using RNA-Rx, we probed S2P-RNA Pol II and H2Bub chromatin enrichment at genes composing the seven DREM modules. Paths A’ to C’ displayed stronger enrichments of these transcription hallmarks in light than in dark. This was also true when comparing path D’, in line with the idea that a majority of genes are upregulated instead of being stably expressed (**Fig. 4D, Figure S6B**). Interestingly, genes in path E’ displayed a mild increase in H2Bub and S2P-RNA Pol II mean levels in the light, indicating that epigenomic data imperfectly match both RNA-seq and RNA-Rx analyses for this intermediate path. In contrast, genes in paths F’ and G’ displayed low markings in the light, especially S2P, in line with their downregulation upon the transition.

One of the striking findings using RNA-Rx was a massive correction for fewer downregulation than previously assumed. This compelled us to investigate whether the transcripts of downregulated genes were particularly unstable. The transcript half-live estimates from recent studies^43–45^ were used to test the intrinsic mRNA stability of genes composing the seven DREM paths. In agreement with the assumption that transcriptional shutdown and mRNA decay both contribute to transcript depletion, mRNAs of the downregulated paths F’-G’ genes displayed a tendency for short half-lives (**Fig. 4E**). Based on studies employing cordycepin (3’-deoxyadenosine) to block mRNA synthesis, the paths A’ and B’ corresponding to highly upregulated genes also tend to encode short-lived transcripts. In contrast, with thousands of moderately upregulated genes (∼8,700), paths C’ and D’ tend to encode transcripts with a long half-life (**Fig. 4E upper panels**). This observation echoes studies in mammals showing that transcripts mediating constitutive cellular processes, such as housekeeping gene mRNAs, tend to be more stable than those of transcription factors and signaling components^46^. A slight increase in transcription rates of transcripts with a long life may therefore contribute substantially to transcriptome size doubling during cotyledon de-etiolation.

Given these observations, we revisited the RNA-Rx-defined co-expression modules to gain an updated understanding of light-regulated biological functions. Upon Gene ontology (GO) analysis, path D’ with discordant trends between standard RNA-seq and RNA-Rx was significantly enriched in genes involved in replication and translation, including most ribosomal protein genes as well as RNA Pol I and RNA Pol III subunits (**Fig. 4F-G, Figure S6C**). Hence, contrary to standard RNA-seq, RNA-Rx identified the upregulation of the replication and translation functions, supporting the known positive effects of light on translation^11,47–51^ and the onset of endoreduplication occurring 24/48 hours after exposure to light^11^. Taken together, these results demonstrate that regulatory networks defined using RNA-Rx better support epigenomic profiling data and are more coherent than standard RNA-seq in terms of biology.

### Identification of stably expressed genes to quantify transcriptome size changes in standard RNA-seq data

Considering that defining an internal reference would open the way to absolute quantification of transcriptome variations in RNA-seq datasets lacking a spike-in and may further reduce technical variability, we looked for genes whose expression patterns mimic the spike-in information. We sorted out the genes displaying moderate/high transcript levels and a high signal-to-noise ratio and ranked these genes based on the likelihood ratio, estimating their ability to recapitulate the spike-in information (**Fig. 5A**; see methods). As detailed in **Figure S7A**, the top 32 genes were selected as our candidate set of stable genes (**Fig. 5B-C**). These genes encode proteins with basic molecular and cellular functions, such as organelle and vesicle trafficking, transcription and RNA metabolism, splicing, ribosome biogenesis, and vacuolar H^+^ ATPase subunits (**Fig. 5B**). Interestingly, 28 of them are poorly translated in response to darkness or hypoxia^48,50,52^ indicating that genes stably expressed during cotyledon de-etiolation may predominantly be regulated post-transcriptionally.

**Figure 5.**
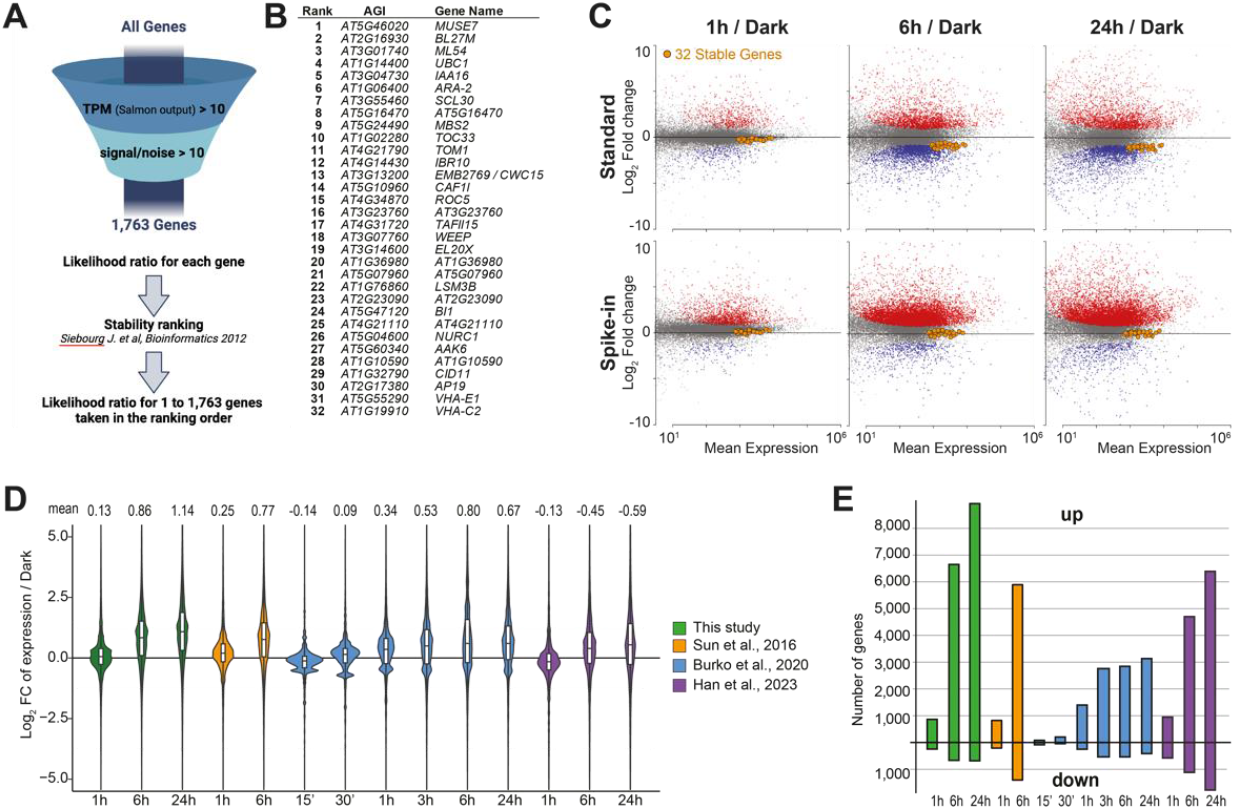
Genes with stable absolute expression recapitulate spike-in information in independent studies of cotyledon de-etiolation. **A.** The pipeline used to identify reference genes with stable absolute expression levels during cotyledon de-etiolation led to the selection of 32 genes (see Methods for details). **B.** Identity of the selected stable genes. **C.** MA-plot analyses after Standard RNA-seq or Spike-in RNA-Rx highlighting the 32 stable genes (orange). **D.** Differential gene expression in our study and in 3 published datasets^8,9,13^ using the set of 32 stable genes as an internal reference for normalization. In each analysis, the dark time point was used as reference. **E.** Number of differentially expressed genes (q-value < 0.01 for a |Log2 FC| > 0.5) using the 32 reference genes as an internal control for normalization of our RNA-seq data and of the same published datasets as in (D).

We tested the capacity of the stable gene set to provide equivalent information to exogenous spike-in. Renormalization of the de-etiolation RNA-seq time series on this gene set as reference expectedly resulted in differential expression changes highly correlated to those obtained with Drosophila spike-in as reference (R^2^ ≥ 0.89 at all time points; **Figure S7C**) and a comparable doubling of transcriptome size at 24 hours (**Fig. 5D-E and Figure S7B**).

We tested whether the selected set of stable genes could be used to faithfully detect transcriptome size changes in RNA-seq data obtained without spike-in. We remapped and normalized three publicly available transcriptomic datasets of cotyledon deetiolation (**Figure S8A**; Han et al. 2023^8^; Sun et al. 2016^9^; Burko et al. 2020^13^) using the set of 32 genes as an internal reference. Remarkably, the distribution of transcript levels and differential expression changes consistently detected a similar augmentation of transcriptome size in the four independent laboratory setups (**Fig. 5D and Figure S8B-D**). In our RNA-seq time series, transcriptome size variations were nonetheless consistently lower at the 1-hour and 6-hour time points using the stable gene set than the Drosophila spike-in (**Figure S7B-C**), indicating that normalization with stable genes may slightly underestimate global variations. Accordingly, the number of upregulated genes was lower in the four renormalized datasets than using a proper spike-in normalization, especially in the study by Burko et al. that displays lower number of replicates and smaller library sizes (**Fig. 5E and Figure S8D**). Yet, with an average trend of 84% upregulated versus 16% downregulated genes at 6 hours, normalization using the set of 32 stable genes reproducibly detected clear trends toward gene upregulation in all four studies (**Fig. 5E**). Our data renormalization technique thus enabled the robust detection of an overlooked transcriptome size increase in all tested independent studies.

## Discussion

*Arabidopsis* photomorphogenesis is a developmental transition that profoundly reshapes the transcriptional program^1–3,15,53^. In previous studies, we proposed that nuclei of skotomorphogenic cotyledons display a quiescent transcriptional state^17,19^. Here, we unveiled that cotyledon photomorphogenesis involves a rapid increase of the transcriptional regime linked to a quasi-exclusive trend of gene upregulation and transcriptome expansion. The existence of hyper- and hypotranscriptional states in plants has long been overlooked, and the capacity of eukaryotic cells to adjust their transcriptional regime to developmental or external signals is just starting to emerge^24,54^. The biological significance and underlying mechanisms of hyper-transcription are best understood in Embryonic Stem Cells (ESCs), tumors, and multiple progenitor cell lineages, which all over-express c-Myc^20,23,55–57^. Vice versa, hypotranscription has been linked to specific metabolic states, *e.g.* during yeast cell quiescence^58–60^ and mouse blastocyst diapause^54,61^. In plants, hypotranscription has been suggested in spore mother cells^62^, in the egg cell and resulting embryo^63^, and in dry seeds^64,65^. Global variations of RNA Pol II activity were also found to follow the changes in genome content upon endoreduplication or polyploidization^26,27^ as well as during the cell cycle with tissue-specific effects^31^. Our study further unveiled that environmentally-induced plant cellular transitions can involve changes in the transcriptional regime at the genome scale and additionally decrypted their impact on transcriptome dynamics.

Determining the impact of higher-order control of transcription on individual genes’ expression and transcriptome size requires absolute normalization approaches, a bottleneck that we addressed using an exogenous reference. Previous studies assessing the effect of plant polyploidization on transcriptome size have considered spike-in strategies for a normalization per biomass, cell, or genome content^24,28–30^. A ‘per-leaf normalization’ also recently unveiled that sorghum leaves display higher average transcript levels in the evening than in the morning^66^. Given that cotyledon de-etiolation initiates independently of mitosis and endoreduplication, here we were able to reach an absolute normalization averaged per cell, which was essential to detect and quantify a global increase in mRNA cell content during cotyledon photomorphogenesis. The doubling in transcriptome size identified during the first six hours cannot be explained by endoreduplication, negligible during this short time, but may more directly result from the 2-fold increase of active RNA Pol II amounts per cotyledon nucleus that we detected *in vivo*. This likely initiates before the onset of endocycles to sustain mRNA and protein production upon full adaptation to light. Light sensing indeed strongly enhances global translation in plant cells^53^, a function evidenced in our RNA-Rx data by the upregulation of genes involved in protein synthesis. Studies employing exogenous spike-in RNA as a normalization reference to quantify polysome-bound mRNAs reported an overall increase in translation four hours after the induction of Arabidopsis photomorphogenesis^47,48,50^.

Translation and transcription are both energetically costly^67^, hence attenuation of these two basic cellular activities plausibly constitutes an adaptation to the low metabolic status upon prolonged growth in darkness. Etiolated development relies on limited seed resources that, in the absence of photosynthesis, can only last a few days^4^. Yet, the control of translation is surprisingly independent of photosynthesis but relies on an auxin-TOR signaling path acting downstream of COP1^50^. Translation inhibition partially relies on mRNA sequestration in cytosolic p-bodies that are particularly abundant in darkness^48^. Interestingly, most of the stably expressed genes identified here using an RNA-Rx approach correspond to genes regulated at the translational rather than transcriptional level. Considering the hypotranscriptional state of etiolated cotyledons, we hypothesize that transcripts stored within p-bodies are rapidly available for translation to enable the seedling to initiate photomorphogenesis while reaching a full transcription competency.

Regulation of translation may itself impact the transcriptional regime, for example by sustaining the production of unstable chromatin modifiers as achieved by the TOR pathway in mouse embryonic stem cells^68^. In etiolated cotyledons, COP1/auxin/TOR-mediated translation control may similarly attenuate the transcriptional regime through epigenome adaptations. For instance, DET1 regulates the stability of a histone H2Bub deubiquitination module, and targeted degradation of chromatin modifiers has been proposed to contribute to adjusting the epigenome landscape to a low transcriptional status in darkness^19^. Chromatin regulatory processes are at the nexus of many light-driven signaling pathways and their possible implication in energy-saving mechanisms remains to be tested.

In this study, absolute quantification of transcript levels resulted in drastically revisiting former views on transcription factors’ activity at thousands of light-regulated genes. This further shed light on higher-order regulatory levels that encompass the whole transcriptome. Disentangling the contribution of genome-scale transcription intensification from sequence-specific control at individual genes will, therefore, be important to decrypt the molecular mechanisms modulating transcriptome size augmentation. A second layer of complexity is the existence of cell-to-cell variability in gene transcriptional responses to environmental changes identified through *in situ* imaging of transcription reporter transgenes^69,70^ and RNA FISH^71^. These case studies identified that organ-level transcription regulation combines two patterns: an ‘analogic’ mode involving gradual adjustments of gene transcription across all cells and a ‘digital’ mode in which genes are switched ON and OFF in distinct cells^69,71,72^. Accordingly, single-cell and single-nucleus RNA-seq^8,73,74^ combined with ad hoc absolute normalization methods may allow disentangling the contribution of analog and digital regulation modes to transcriptome size changes.

## Methods

### Plant material

All seed lots were *Arabidopsis thaliana* Col-*0* wild-type plants, except for mintbody microscopy that used a *pRPS5a:Ser2P-mintbodyIntF2A-H2B-mRuby* line^31^ and a *CYCB1;1::DB-GUS* reporter line^75^.

### Growth conditions

Seeds were surface sterilized and sown on half-strength Murashige and Skoog media (Duchefa Biochemie, The Netherlands) with 1% (w/v) Phytoagar, pH 5.7. After 3 days of stratification at 4°C in darkness, seeds were exposed to light for 4-6 h to synchronize germination. Plants were either grown under light conditions for 5 days (with a 16 h-light 8 h-dark photoperiod) or under three layers of aluminum foil for 5 days before being harvested (Light and Darks samples, respectively) or exposed to 1 h, 6 h, or 24 h of continuous light (80 µmolm-2m-1 irradiation from cool white fluorescent tubes (Philips F25T8/TL841 (Hg)) in a Percival CU36L5 growth chamber. Dark samples were collected under a dim green light.

### RNA extraction and library preparation

The light-exposed or Dark samples were collected in 80% acetone at -20°C before vacuum infiltration twice for 5 min under white or dim green light, respectively. Unless stated otherwise, 200 cotyledons were dissected in 100% ethanol under a dissection scope, and frozen in liquid nitrogen before grinding with a Tissue Lyser (Qiagen). RNA was extracted with the microRNeasy kit (Qiagen). Five million *Drosophila melanogaster* S2 cells were resuspended in 7 mL of RLT buffer containing ß-mercaptoethanol. For this, cotyledons were ground in liquid nitrogen and resuspended in 700 µL of the RLT buffer containing the S2 cell lysate by vortexing and repeated resuspension through a 20-gauge (0.9 mm) needle attached to a sterile plastic syringe. Combined purification of *A. thaliana* and *D. melanogaster* RNA was then conducted according to the manufacturer’s instructions. Sequencing libraries were prepared using the Illumina TruSeq Stranded mRNA kit with polyA selection. Paired-end sequencing was performed on a DNBSEQ-G400 platform by BGI (Hong Kong) with a read length of 100 base pairs.

### Read mapping

After trimming, reads were mapped using Salmon^76^ (version 1.9.0) on a merged reference transcriptome composed of the AtRTD3^77^ Arabidopsis and the dmel_r6.48 Drosophila transcriptomes and using the Arabidopsis (TAIR10) and Drosophila (dmel_r6.48) genome as a decoy sequence. Salmon options to correct for sequence and GC bias were used and 100 Gibbs inferential replicates of the count assignment were sampled with a thinning factor of 100. The read counts were obtained from the median of those inferential replicates via the tximport^78^ R package (version 1.30.0, R version 4.3.2) with the ‘scaledTPM’ option. For the rest of the processing steps, only the transcripts annotated as mRNAs in AtRTD3^77^ and having at least 1 read count in 1 of the samples were considered.

### Normalization and differential analysis for Standard RNA-seq

Differential analysis was performed using DESeq2^34^ (version 1.42.0, R version 4.3.2) using read counts on Arabidopsis genes as input. The DESeq2 design matrix included an additive ‘batch’ effect thus taking into consideration the fact that 5 biological replicates were obtained in two series. Genes were considered differentially expressed at a q-value threshold of 0.01 for an absolute Log_2_ fold change greater (or lesser for stably expressed genes) than 0.5 by setting the ‘lfcThreshold’ option in DESeq2 ‘results’ function to 0.5. Normalized transcript levels were obtained from the DESeq2 normalized read counts divided by gene length and scaled to arbitrarily set the dark sample’s median to 1 for each analysis. For scatter plot comparisons of datasets, the Log_2_ fold change determined using DESeq2 was shrunk with the ashr^79^ package (version 2.2.63, R version 4.3.2) using a mixture of Normal prior, no point mass at 0 and the ‘estimate’ option for the prior mode.

### Normalization and differential analysis for Spike-in RNA-Rx

To mitigate read count dispersion due to spike-in variability (**Supplemental Figure 3C)**, we used the CATE^80^ package (version 1.1.1, R version 4.3.2). First, each sample was normalized using the DESeq2 median of ratios computed on the Drosophila gene read counts. The Log2 of the obtained Arabidopsis genes’ normalized read counts (using a 0.5 offset) and the same design matrix used for Standard RNA-seq (including a ‘batch’ effect) was used as input for CATE to adjust for the principal latent confounder (‘r’ set to 1 in ‘cate’ function). The ‘cate’ function was run in ‘negative controls’ mode using Drosophila genes with at least 5 reads in all samples. The obtained matrix of corrected counts was used for DESeq2 differential analysis with the size factor calculated from the median of ratios on Drosophila genes corrected counts and with the already described design matrix including an additive ‘batch’ effect. Differentially expressed genes, normalized transcript levels, and scatter plots were retrieved in the same way as for Standard RNA-seq.

### DREM analysis

Log_2_ fold change of expression from Standard RNA-seq and Spike-in RNA-Rx differential analyses were used as inputs using the same DREM parameters as in Bourbousse et al. 2018^81^.

### Selection of stably expressed genes

Plastid, mitochondrial, AtRTD3-newly identified transcripts, and AtRTD3 annotations overlapping multiple genes were excluded. Only genes whose TPM median and signal-to-noise ratio minimum across samples were both greater than 10 were considered. TPM values were retrieved directly from Salmon^76^ (version 1.9.0) using tximport^78^. The signal-to-noise ratios were calculated as the median divided by the square root of the variance of the gene counts across the inferential replicates for each sample. For each of the 1,763 remaining genes, size factors were calculated for each sample (similar to the DESeq2’s median of ratios method, i.e., as the ratio of the sample’s gene counts over the geometric mean of gene counts across all samples) using CATE corrected counts. We compared the size factors obtained for each of these 1,763 genes with those obtained with the Drosophila spike-in genes using the coefficients and their standard error derived from the linear regression of log-transformed size factors on the sample’s design matrix. Specifically, for each candidate gene, we calculated (1) the ‘absolute deviation’ as the maximum of the absolute difference between the candidate gene’s coefficients and the Drosophila spike-in coefficients, (2) the ‘variability’ as the maximum of the candidate gene’s coefficient’s standard deviations and (3) the ‘likelihood ratio’ as the minimum of the ratio of the spike-in model coefficients’ likelihoods in the gene model over the spike-in model. For those 3 measures, the coefficient corresponding to the intercept was ignored. Genes were ranked using the likelihood ratio as a ranking score using a method similar to the stability ranking procedure described in Siebourg et al. 2012^82^ to increase robustness. 100,000 rankings of the candidate genes were computed from count matrices built by sampling genes count independently from Salmon inferential replicates of each sample, and the final ranking was obtained with a probability threshold^82^ π = 0.95. The counts of each inferential replicate were also corrected separately using the CATE package as described above. According to this final ranking, the ‘absolute deviation’, ‘variability’, and ‘likelihood ratio’ were calculated this time using an increasing number of candidate genes. The ‘likelihood ratio’ is stable across the first 100 genes, yet we defined the final set of stable genes as the first 32 ranked genes given the presence of local maximum at this position (**Figure S7A**).

### Normalization and differential analysis using stable genes as endogenous references

Normalization, differential analysis, and absolute transcript levels computation with endogenous reference were conducted similarly to the Drosophila spike-in method described above by replacing the Drosophila genes with the 32 stable genes but without implementing a ‘batch’ effect correction.

### ChIP-seq

Dark and light-grown seedlings were fixed in 1% formaldehyde under dim green or white ambient light, respectively. A fixed number of 400 cotyledons were dissected, frozen, and ground. For S2P RNA Pol II ChIP, a second crosslinking step was performed and chromatin was extracted as in Bourbousse et al. 2018^81^, sheared using a Covaris sonicator, and immunoprecipitated using Abcam Ab5095 antibody. For H2Bub ChIP, a chromatin spike-in was included and immunoprecipitation was conducted as in Nassrallah et al. 2018^19^ using the Medimabs MM-0029-P antibody. Libraries were prepared using NEBNext® Ultra™ II DNA Library Prep Kit for Illumina® (NEB, E7645). Paired-end sequencing was performed on a DNBSEQ-G400 platform by BGI (Hong Kong). ChIP-seq bioinformatic analyses were conducted following the pipeline presented in the accompanying GitHub resource.

### RT-qPCR

Reverse transcription was performed using the High Capacity cDNA Reverse Transcription Kit (Applied Biosystem) using random hexamers followed by quantitative PCR using the SYBR Green I Master Mix (Roche) with the oligonucleotides listed in **Supplemental Data 8**. PCR analyses of *AtRS31* pre-mRNA were conducted as in Herz et al. 2019^33^. Absence of DNA contamination was checked for all RNA samples by amplification without the RT enzyme.

### Immunocytometry of RNA Pol II

Seedlings were fixed under safe green light in Galbraith buffer (20 mM MOPS, 45 mM MgCl2, 30 mM sodium citrate, 0.3% Triton X-100) containing 1% formaldehyde under vacuum for 20 min then releasing vacuum and leaving the seedlings in the fixative without vacuum for 20 min. The fixed seedlings were rinsed three times with water and transferred to Galbraith buffer containing 5 mM sodium metabisulfite (chopping buffer) on ice. A fixed number of 160 etiolated or 80 de-etiolated cotyledons were dissected in a small drop of chopping buffer and then chopped with a razor blade on glass Petri dishes. Upon sample collection in 1.8 mL of chopping buffer and filtration through a pre-wetted 30 μm nylon mesh filter, the nuclei were pelleted by centrifugation at 1,500 *g* for 5 min at 4°C and resuspended in 100 μL of antibody buffer (30 mM MOPS, 22 mM MgCl2, 15 mM sodium citrate, 300 mM sorbitol, 0.5% BSA, 0.3% Triton X-100) containing the Abcam ab5095 and Abcam ab817 primary antibodies at a 1:500 dilution. Upon incubation overnight at 4°C the nuclei were washed with 1.9 mL of antibody buffer (devoid of antibody), pelleted at 1,500 *g* for 5 min at 4°C, and resuspended in 150 μL of antibody buffer containing Alexa488 coupled anti-mouse (Thermo Fisher A11001) and Alexa568 coupled anti-rabbit (Thermo Fisher A11011) secondary antibodies at 1:400 dilution. Upon incubation for 2 h at ambient temperature in darkness, the nuclei were washed with 1.85 mL of antibody buffer, pelleted at 1,500 g for 5 min at 4°C, and resuspended in 200 μL of antibody buffer with 5 μg/mL of DAPI. The immunolabeled nuclei were acquired with a Cytoflex S bench-top cytometer (Beckman-Coulter) in a 96-well plate, driven by cytExpert 2.5. Data analysis was made with the Kaluza 2.1 software (Beckman-Coulter). Nuclei were recognized and classified by ploidy level according to the DAPI staining for gDNA. Doublet nuclei were excluded using the cytogram of DAPI-Area versus DAPI-Heights signals. The negative population for immunostaining was defined on the Alexa Fluor 488 and 568 background signals from a control sample lacking the primary antibodies. From this, a NOT gate was applied to all fully-labeled samples. Finally, a customized parameter was used to measure the fluorescence ratio of Alexa488 and Alexa568 for each nucleus from each ploidy level. For each ploidy level, gated data were extracted from Kaluza in CSV file format for further R analysis.For statistical analysis, we employed a linear model on the log-transformed response variable (S2P/NP ratio, S2P signal, or NP signal) to account for heteroscedasticity and stabilize variance across treatments. The model was fitted using the following formula: ∼Condition * Ploidy + Experiment, where “Condition” refers to the light condition and “Experiment” to the experimental batch (one for replicate 1 and 2 and another one for replicate 3). Following model fitting, we used specific contrasts to test hypotheses using a t-test with the “HC3” heteroscedasticity consistent covariance matrix estimator^83^.

### Ploidy measurements

Cotyledons were chopped in Galbraith buffer with 0.3% Triton X-100. DAPI was added to a final concentration of 20 µg/mL^-1^. Nuclei were counted in each sample according to their ploidy level determined by fluorescence intensity using a Cytoflex bench-top cytometer (Beckman-Coulter). For statistical analysis, the ploidy count data of each sample were first transformed with the additive log-ratio transform (ALR) using Bayesian-multiplicative approach for count 0 imputation considering a Bayes-Laplace prior^84^.The p-values were obtained using an unequal variance Hotelling test.

### GUS staining

*CYCB1;1::DB-GUS* seedlings were fixed in 80% cold acetone before incubation with 2mM X-Gluc staining solution (50 mM NaPO4 pH 7.2, 0.2% Triton X-100) at 37°C. After washes in 70% EtOH, seedlings were further discolored using chloral hydrate (2.5 g.mL^-1^ in 30% glycerol).

### Mintbody sample preparation, image acquisition, and 3D segmentation

Live seedlings were mounted in Immersol W 2010 (Carl Zeiss Microscopy, LLC, USA). Living nuclei from mesophyll cotyledon cells were imaged with a Leica TCS SP8-MP (Leica, Germany) equipped with a resonant scanner (8kHz) with a 63x water objective (HC PL APO 63x/1.20 W motCORR CS2 1.2 NA). EGFP and mRuby were excited with 960 nm and 1040 nm lasers (Insight DS+ Dual (680-1300 nm & 1041nm) ultrafast NIR laser for multiphoton excitation), respectively. Fluorescence was detected using a non-descanned super-sensitive photon-counting hybrid detector (HyD), operated in photon-counting mode with 8× frame accumulation. z-Stacks were composed of 8-bit images acquired with a resolution of 568×568 pixels, and the voxel size was 0.0903 x 0.0903 x 0.0901 μm. All images were deconvolved with Huygens Professional version [23.04] (Scientific Volume Imaging, The Netherlands) and segmented with Imaris (bitplane, Switzerland) using the Surface creation tool in manual contouring mode for the nucleolus and ML Pixel Classifier mode for the nucleus. For each nucleus, channel signals (EGFP, mRuby, and their ratio) were exported and used to compute sums and means.

## Data availability

Raw sequencing data and processed files for RNA-seq and ChIP-seq, and source data will be provided upon peer-reviewed publication.

## Code availability

ChIP-seq scripts are available at https://github.com/vidal-adrien/ChIP-Rx-Pipeline-Pub and RNA-seq scripts are available upon request.

## Acknowledgments

The authors are grateful to Chris Bowler (IBENS, France) for constant support, Vincent Colot (IBENS) and Etienne Delannoy (IPS2, France) for constructive feedback, Pierre Vincens and the Bioinformatics platform (IBENS) for technical support, Frédérique Perronet (IBPS, Sorbonne Université, France) for sharing Drosophila S2 cells, Ouardia Ait-Mohamed (formerly IBENS) for help with housekeeping genes’ analyses, to Dr Mio K. Shibuta (Yamagata University, Japan) and Prof Sachihiro Matsunaga (Tokyo University, Japan) for kindly providing the *pRPS5a:Ser2P-mintbodyIntF2A-H2B-mRuby* line. We thank Dr Emmanuelle S. Botté for her editorial advice.

## Contributions

CBo and FB conceived and coordinated the study. RNA-seq experiments were performed by CBo and LW; ChIP-seq experiments by CBo, EA, and DC; RNA Pol II immunocytometry by MDa with the contribution of MB; RNA pol II imaging of nuclei and analysis by FMM and CBa. GS developed the RNA-Rx analysis pipeline and analyzed the RNA-seq data. CBo analyzed the ChIP-seq data. AV developed the GitHub repository. CBo prepared the figures. FB and CBo wrote the manuscript. All authors had the opportunity to edit the manuscript and have approved it.

## Supplementary Figures

**Figure S1.**
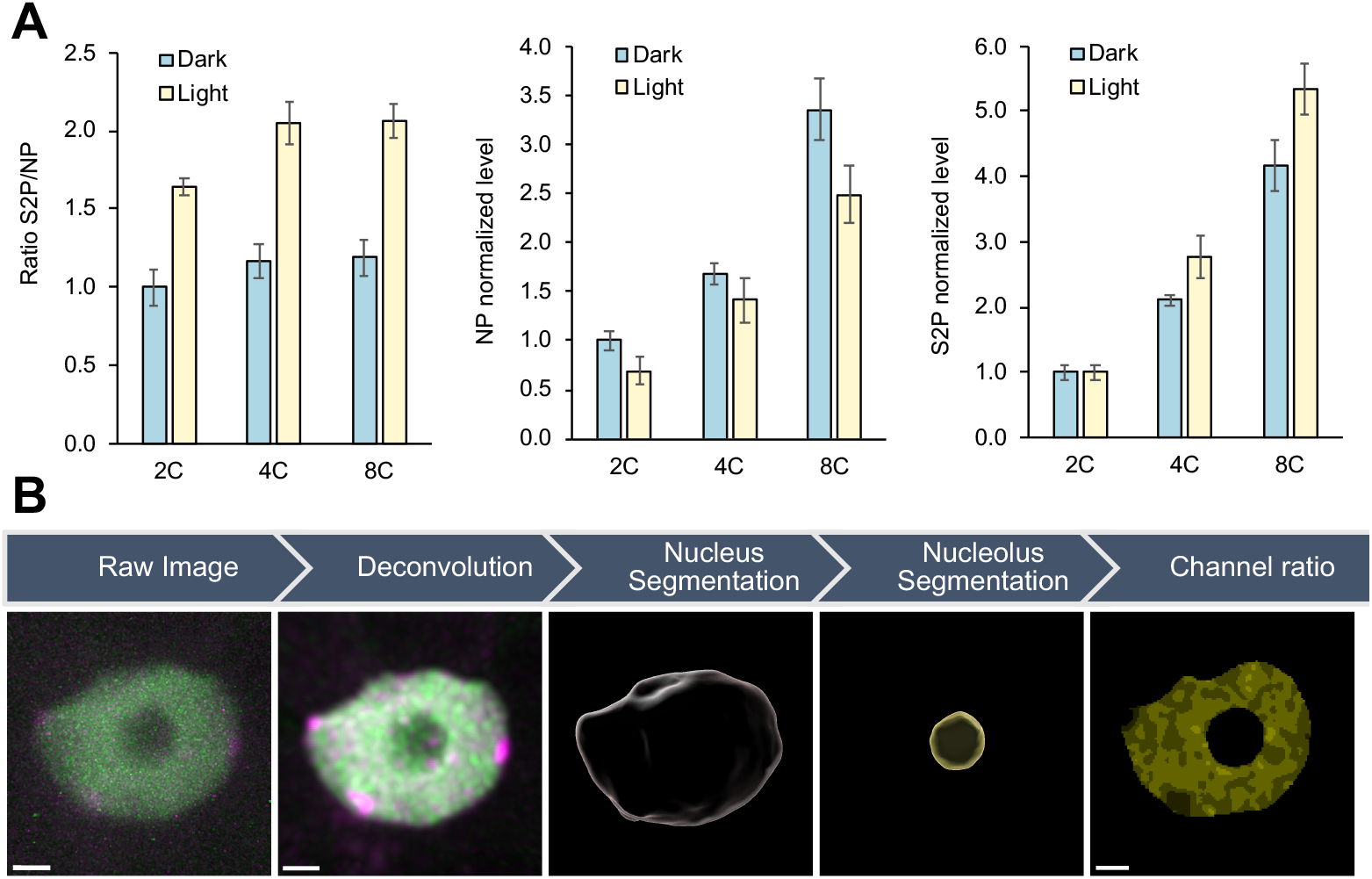
**A.** Mean and standard deviation between the three biological replicates of the immunolabeling signal of RNA Pol II inactive (NP) and elongating (S2P) forms, respectively, for each ploidy class. Fluorescent signals of immunolabelled RNA Pol II forms in cotyledon nuclei correspond to the flow cytometry experiment of Figure 1B. The mean of 2C Dark samples was arbitrarily set to 1. **B.** Analysis workflow of live RNA Pol II mintbody signal acquisition and image 3D segmentation. (i) raw image of a 3D projection from multiphoton live image reporting S2P mintbody-GFP (active RNA Pol ll) and H2B-mRuby (chromatin); (ii) same image after deconvolution; (iii) the nucleus is segmented through machine learning; (iv) the nucleolus is segmented manually; (v) sub-nuclear distribution of S2P RNA Pol ll local enrichment given by the ratio between green and magenta channels in the nucleus excluding all pixels corresponding to the nucleolus. Source data are provided as in Source Data 1.

**Figure S2.**
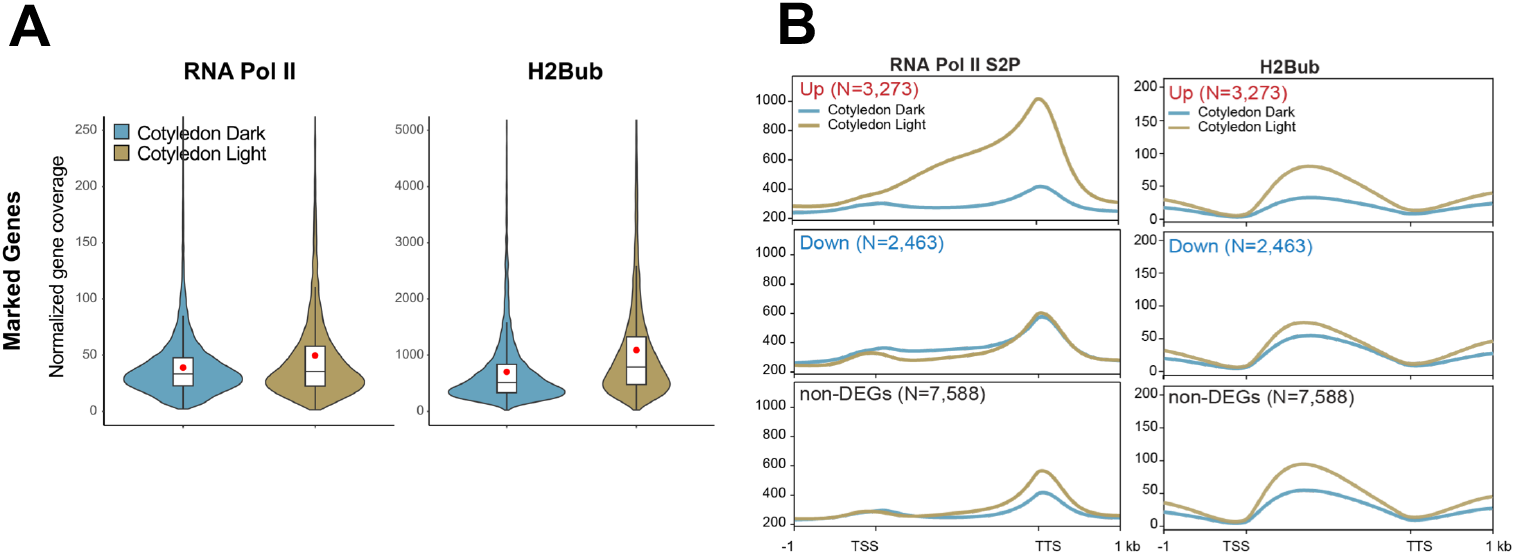
Violin plots showing the distribution of S2P RNA Pol II and H2Bub enrichment at marked genes, corresponding to the ChIP-seq data in Figure 1E. **B.** ChIP-seq meta-profiles of S2P RNA Pol II and H2Bub at up-regulated and down-regulated genes (q-value < 0.01 for a |Log_2_ FC| > 0.5), and non-differentially expressed genes, corresponding to Figure 2C.

**Supplemental Figure 3.**
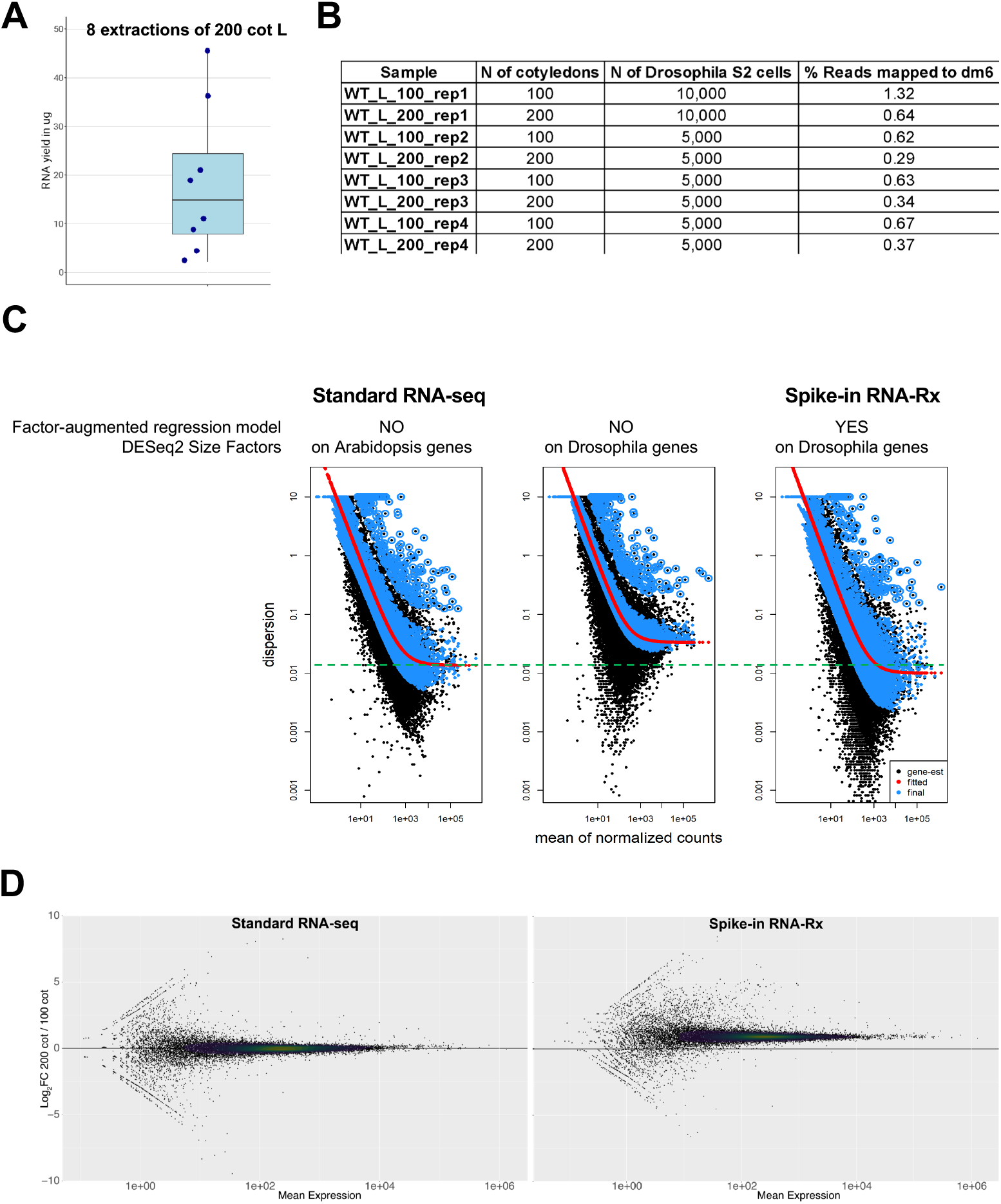
**A.** RNA yield (µg) obtained in 8 independent extractions of 200 cotyledons from light-grown seedlings. **B.** Number of cotyledons and Drosophila S2 cells used in the 4 replicates and the percentage of Drosophila reads retrieved upon sequencing of the mixed Arabidopsis/Drosophila samples for the “100 cot” and “200 cot” samples, referring to Figure 1D-F. **C.** Dispersion pots obtained using standard RNA-seq with DESeq2 default settings, with DESeq2 using the size factors estimated from the Drosophila transcripts and using the modified RNA-Rx pipeline. The estimate for the expected dispersion value according to gene mean expression levels is displayed as a red curve. Each dot is a gene before (black) and after (blue) Shrinking the values toward the curve. **D.** MA-plots showing the distribution of gene expression Log_2_ FC over mean expression level in Standard RNA-seq and Spike-in RNA-Rx analyses.

**Supplemental Figure 4.**
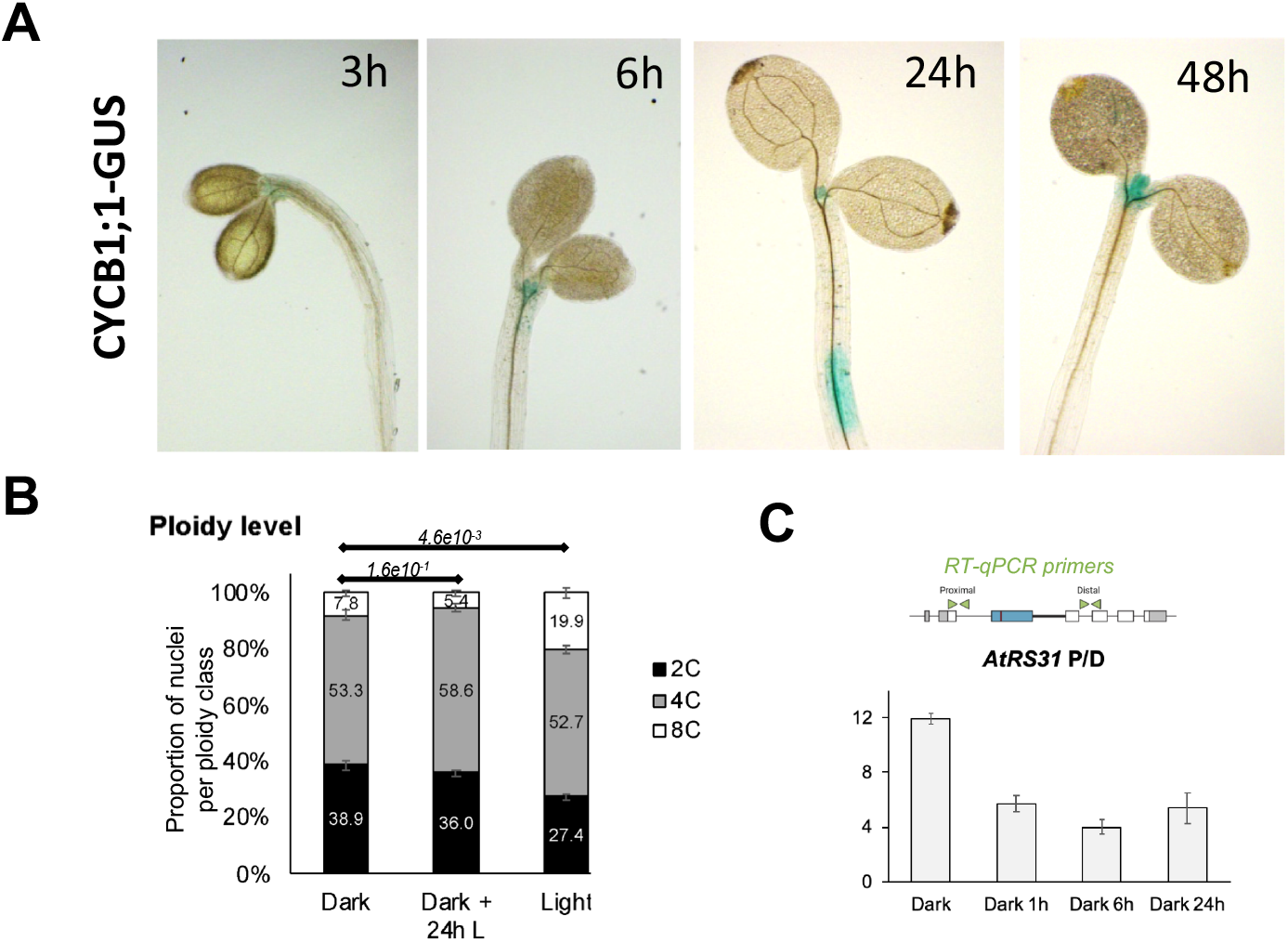
**A.** GUS staining of *CYCB1;1::DB-GUS* seedlings grown for 5 days in the dark and exposed to 3h to 48h of light. **B.** Ploidy class repartition of cotyledon nuclei from seedlings grown for 5 days in the dark and exposed to dark for 0 (Dark) or 24 h of Light (Dark + 24 h L), and from 5-day-old light-grown seedlings (Light). The p-values are given according to a Hotelling statistical test. **C.** RNAs from 5-day-old seedlings grown in darkness and exposed to 0, 1, 6, and 24 hours of light were used to calculate P/D ratios from RT-qPCR signals with primer pairs recognizing a proximal and distal position of *AtRS31* pre-mRNA as in Figure 1C. Error bars correspond to the standard deviation between two biological replicates.

**Supplemental Figure 5.**
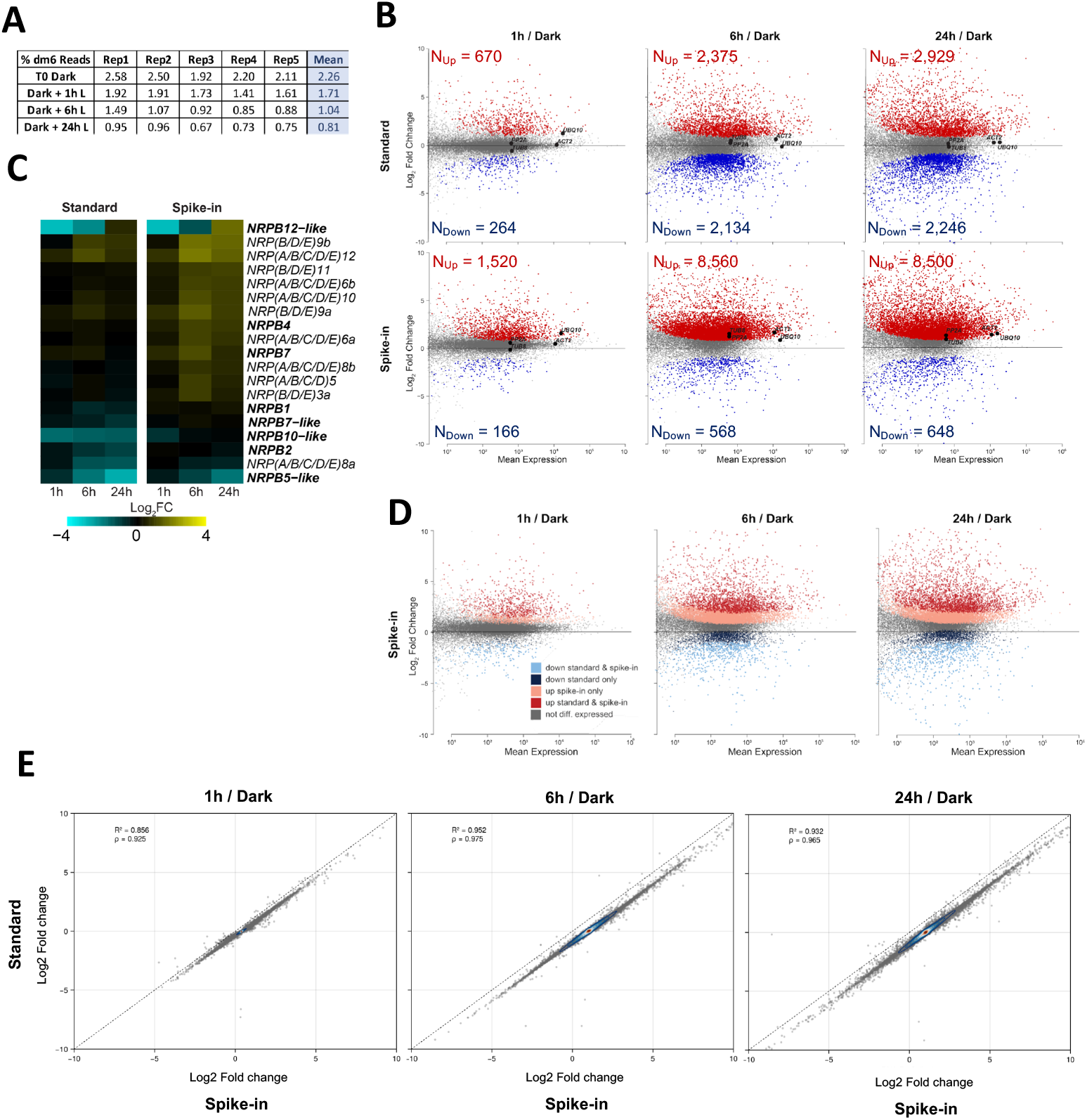
**A.** Percentage of reads mapping to the Drosophila genome in the 20 samples used in Figures 3 to 5. **B.** MA-plots showing the distribution of gene expression Log_2_ FC over mean expression level in Standard RNA-seq and Spike-in RNA-Rx analyses at the 3 time points of light exposure compared with the Dark sample. Red and blue dots represent significantly up- and downregulated genes, respectively (q-value < 0.01 for a |Log_2_ FC| > 0.5 as in Figure 3D). **C.** Gene expression Log_2_ FC at the three time points of light exposure compared to the Dark time point for the genes encoding subunits of RNA Pol II. Genes are ranked according to the Log_2_ FC at 24 h and genes in bold encode RNA Pol II-specific subunits. **D.** Same MA-plots as in B for Spike-in RNA-Rx analyses with dots colored differently if the gene is differentially expressed in Standard RNA-seq, Spike-in RNA-Rx, or both analyses (q-value < 0.01 for a |Log_2_ FC| > 0.5; **Supplemental Data 3**). **E.** Correlation test of gene expression changes during cotyledon de-etiolation in Standard RNA-seq vs Spike-in RNA-Rx using the Dark time point as reference.

**Supplemental Figure 6.**
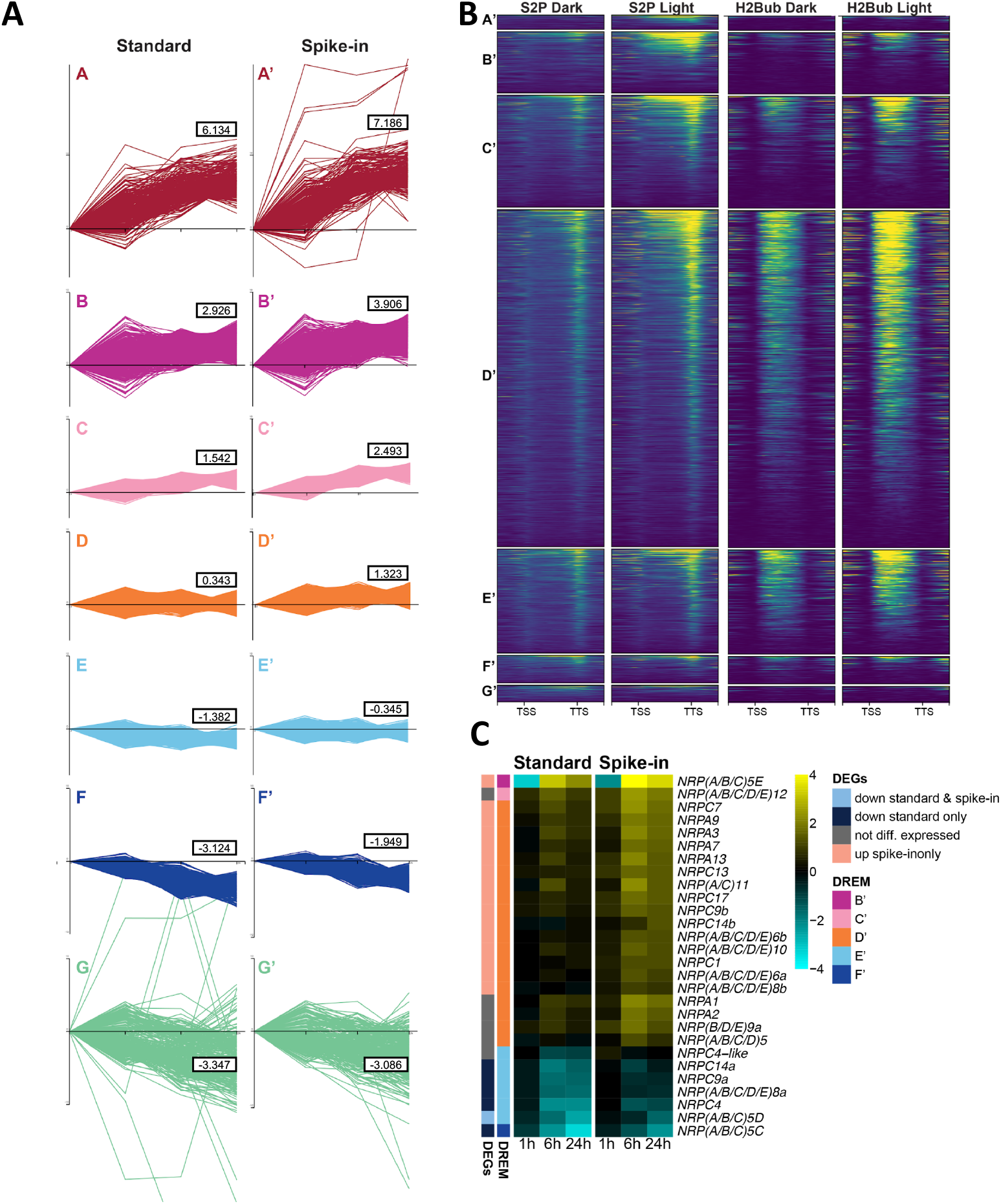
**A.** Time series analysis of individual genes composing the A-G and A’-G’ DREM paths described in Figure 4C. The boxes indicate the mean of the gene expression Log_2_ FC at 24 h. **B.** Heatmaps of ChIP-seq of RNA Pol II S2P form and H2Bub enrichment at genes corresponding to the DREM paths A’-G’. **C.** Expression Log_2_ FC of the genes encoding subunits of RNA Pol I and III. Genes are ranked according to their Log_2_ FC at 24 h and the DREM path and DEG category they belong to are displayed by the color on5the side bars.

**Supplemental Figure 7.**
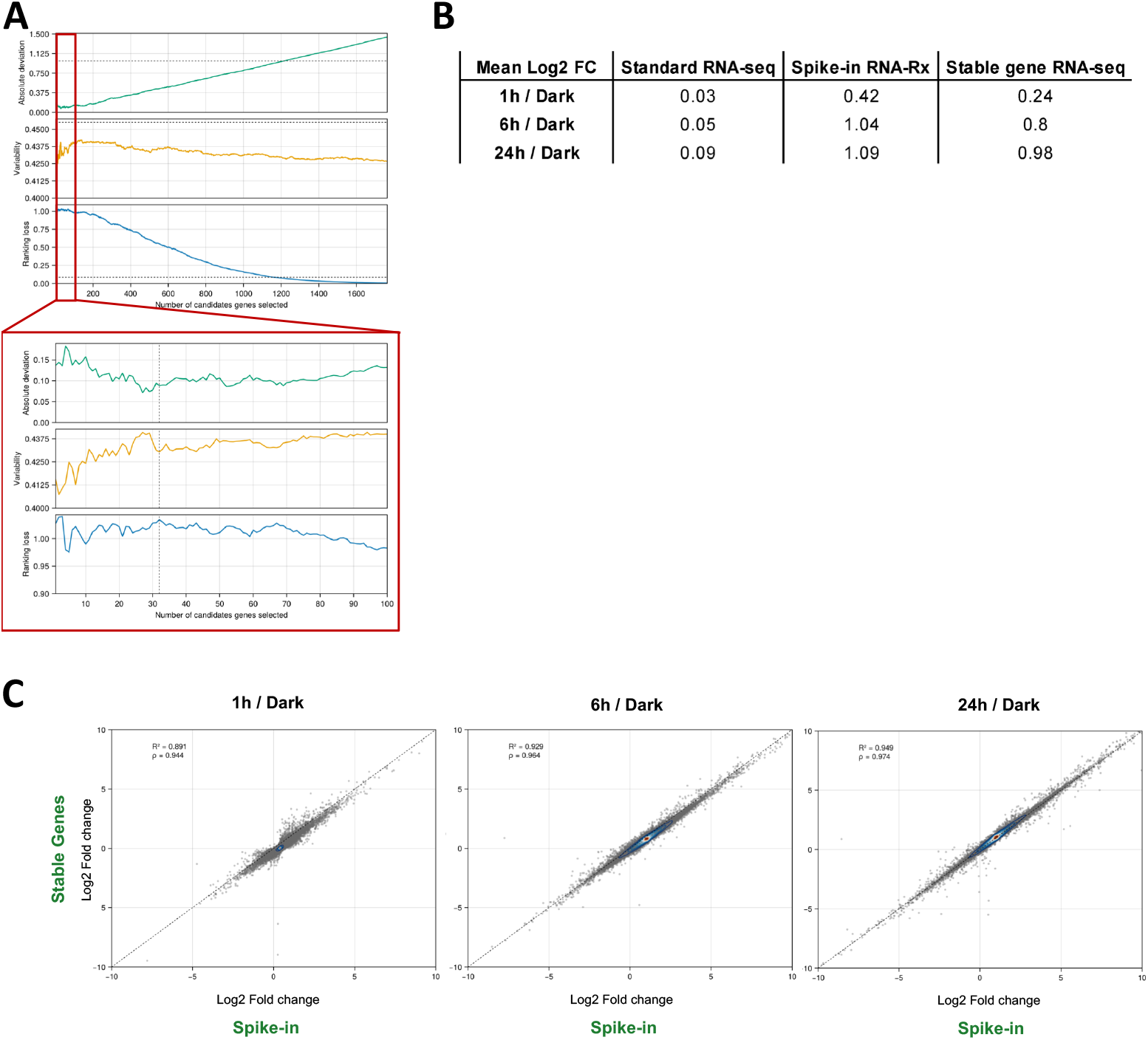
**A.** Trace plot of the absolute deviation, variability, and ranking loss measures (see Methods) obtained with an increasing number (from 1 to 1,763) of the candidate reference genes ranked according to their ability to recapitulate the spike-in information. **B.** Mean gene expression Log_2_ FC at the three time points compared to the Dark time point for each of the three analysis pipelines. **C.** Correlation test of gene expression Log_2_ FC at the indicated time points compared to the Dark time point obtained using the 32 stable genes as an internal reference for data normalization or Spike-in RNA-Rx.

**Figure S8.**
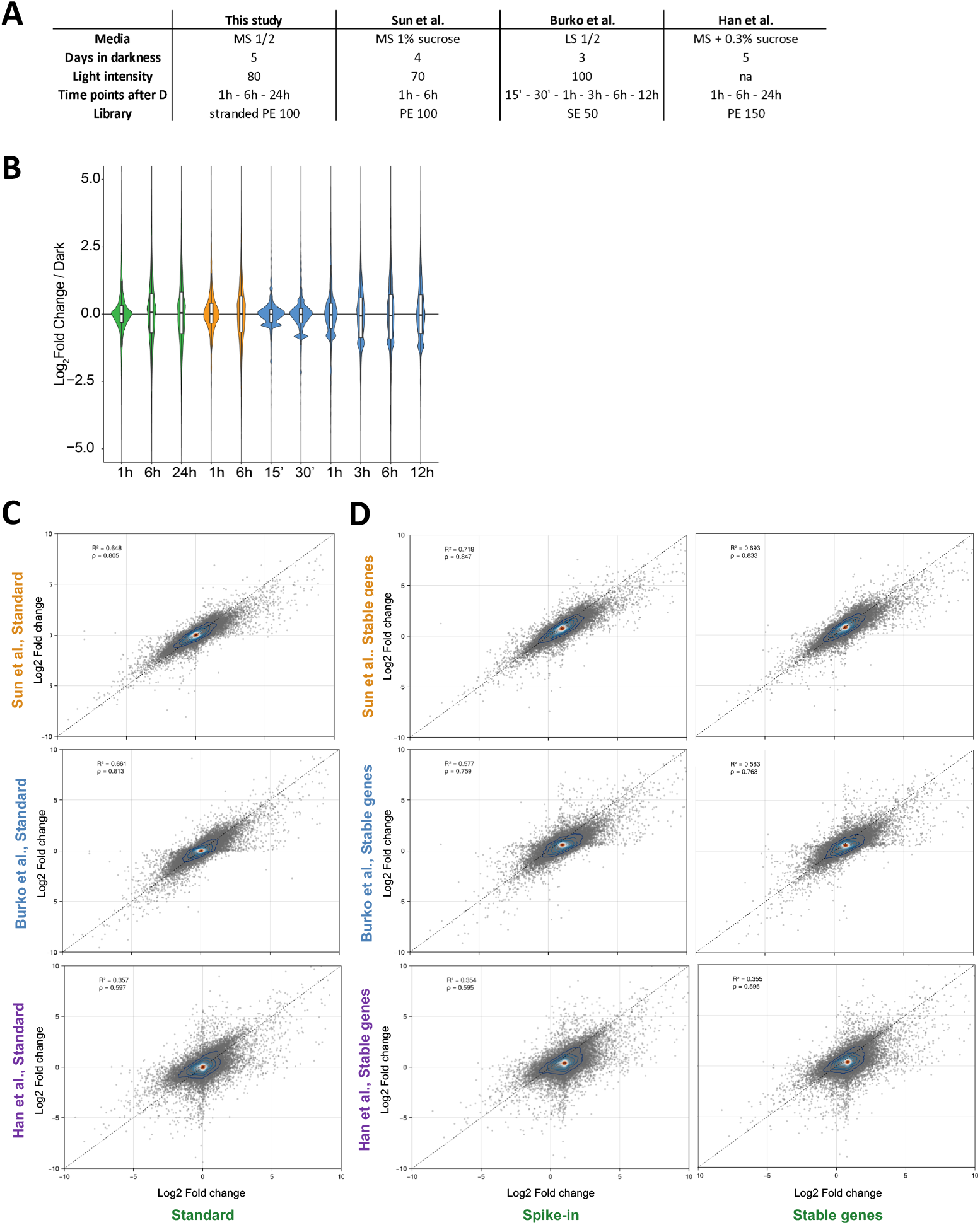
**A.** Table recapitulating the conditions and specificities of the study by Sun et al, Burko et al and Han et al^8,9,13^. **B.** Distribution of gene expression Log_2_ FC between the Dark and other indicated time points in our study and in three published studies^8,9,13^ re-analyzed using our Standard RNA-seq pipeline. **C.** Correlation test of gene expression Log_2_ FC between the three published studies and our dataset, all data being analyzed using the standard RNA-seq analysis pipeline. As shown in (B), datasets for 6 h time-point were available for all four studies. **D.** Same analyses as (C) upon normalization of the formerly published datasets using a set of 32 stable genes as an internal reference compared to our data analyzed using either a Spike-in pipeline or the same set of 32 stable genes for normalization.

